# Histone variant H2A.Z modulates nucleosome dynamics to promote DNA accessibility

**DOI:** 10.1101/2022.08.29.505317

**Authors:** Shuxiang Li, Tiejun Wei, Anna R. Panchenko

## Abstract

Nucleosomes containing the histone variant H2A.Z are important for gene transcription initiation and termination, chromosome segregation and DNA double-strand break repair, among other functions. However, the underlying mechanism of how H2A.Z influences nucleosome stability, dynamics and DNA accessibility remains elusive as experimental and computational evidence are inconclusive. Our modeling efforts of nucleosome stability and dynamics, along with comparisons with experimental data show that the incorporation of H2A.Z results in a substantial decrease of the energy barrier for DNA unwrapping. This leads to spontaneous DNA unwrapping of about forty base pairs in total, enhanced DNA accessibility, nucleosome gapping and histone plasticity, which otherwise is not observed for canonical nucleosomes. We demonstrate that both N- and C-terminal tails of H2A.Z play major roles in these events, whereas H3.3 variant exerts a negligible impact in modulating the DNA end unwrapping. In summary, our results indicate that H2A.Z deposition makes nucleosomes more mobile and DNA more accessible to transcriptional machinery and other chromatin components.

## Introduction

In eukaryotes, genomic DNA is packaged into chromatin using the building blocks called nucleosomes, which are hubs in epigenetic signaling pathways. The nucleosome core particle consists of approximately 147 base pairs of DNA wrapped around an octamer of histone proteins (1). Histones have been historically divided into ‘canonical’ histones that are expressed only during the S-phase of the cell cycle and ‘variants’ that are constitutively expressed during the cell cycle, among other distinguishing features (2). Substitution of canonical histones with histone variants is one of the fundamental ways to dynamically regulate the function of chromatin.

Histone variant H2A.Z belongs to the histone H2A family and is broadly distributed in eukaryotes. H2A.Z is evolutionarily conserved, but human H2A.Z shares only about 60% identity with the canonical H2A (3). H2A.Z predominantly accumulates in the upstream and downstream of the transcription start sites (TSS) and is enriched in a well-defined +1 nucleosome in gene bodies, as well as in enhancer elements of active genes or transcriptionally poised genes (4-7). Given the critical location of H2A.Z-containing nucleosomes in genome and chromatin, one might think that these nucleosomes have major roles in transcription. Indeed, the H2A.Z variant is well known for its functions in transcription initiation and termination (8-10), but there is no unifying view on how H2A.Z incorporation into nucleosomes can modulate the transcription-related processes. On one hand, it has been previously reported that H2A.Z depletion leads to RNA polymerase pausing by the nucleosomal barrier at the transcription start site (11,12). On another hand, the incorporation of H2A.Z may decrease but widen the barrier to transcription (13) and slow the rate of RNAPII pause release (14). In addition, many experimental studies reveal that H2A.Z can accumulate in facultative heterochromatin regions and areas of silenced genes (15,16) and could be involved in heterochromatin boundaries (17) and chromosome segregation (18), facilitate DNA doublestrand break repair (19), and cell cycle progression (20). The functional implications of H2A.Z can perhaps be revealed by studying its physicochemical properties, stability, and dynamics. However, the findings of these studies are also filled with conflicting reports. For example, various experimental works have indicated that the H2A.Z-containing nucleosomes can be less or more stable than the canonical ones (21-23), whereas X-ray crystal structures of canonical and H2A.Z nucleosomes show very similar overall conformations except for the L1 loop region (24). Furthermore, to make the story more complex, it was reported that the stability and dynamics of H2A.Z nucleosomes depended on the nucleosomal context, such as the presence of other histone variants (25,26) or histone posttranslational modifications (PTMs) (27). For example, previous studies have shown that nucleosomes containing the double variants of H3.3 and H2A.Z were enriched at transcription start site (TSS) in human cells and facilitated the access of transcription factors to the packaged DNA (28).

Two major questions remain unanswered in this respect. First, does the H2A.Z variant influence the nucleosome stability, dynamics and DNA unwrapping? Second, are these H2A.Z-influenced events coupled with specific biological functions? To address these intriguing questions, we have performed a computational study by modeling the dynamics and energetics of human heterotypic and homotypic nucleosome systems containing H2A.Z variants (with and without H3.3) and canonical histones. The results, corroborated in multiple six-microsecond long runs, show that incorporation of the H2A.Z variant enhances nucleosomal DNA unwrapping, nucleosome gaping and histone dynamics. Our study also indicates that the C-terminal and N-terminal tails are largely responsible for the altered H2A.Z nucleosome dynamics.

## Results

### H2A.Z deposition facilitates the DNA unwrapping

To investigate the role of H2A.Z in nucleosome dynamics, we first constructed two homotypic nucleosome systems, containing canonical H2A (NUC_H2A/H2A_, Figure 1a) and H2A.Z variant (NUC_H2A.Z/H2A.Z_), and carried out three independent 6 µs long all-atom molecular dynamics (MD) simulations for each system (see Supplementary Figure S1 and Supplementary Table S1 for simulation setup details). We used 187-bp of the native DNA sequence, 147 bp of nucleosomal sequence and two linker DNA segments of 20 bp long each. First, as shown in Figures 1d and 1f, in both canonical and variant systems one DNA end unwraps, while another end stays more bound to a histone octamer. Indeed, previous single-molecule FRET experiments and MD simulations have demonstrated that the DNA unwrapping occurred asymmetrically and DNA unwrapping at one side stabilized the DNA end on the other side (29-31). Moreover, we observe the difference in terms of the degree of unwrapping for both DNA ends that can be related to the asymmetry of our nucleosomal DNA sequence with respect to the dyad position. Second, we find that the overall DNA unwrapping from both ends is significantly enhanced in H2A.Z (up to 40 bps unwrapped in total from both ends) compared to canonical H2A nucleosomes (up to 18 bps from both ends) (Figure 1b). Previous microsecond long simulations and quantitative modeling showed that canonical nucleosomes exhibited spontaneous DNA unwrapping only for systems without histone tails (30-32). The roles of histone tails in DNA unwrapping will be discussed in the next sections.

**Figure 1:**
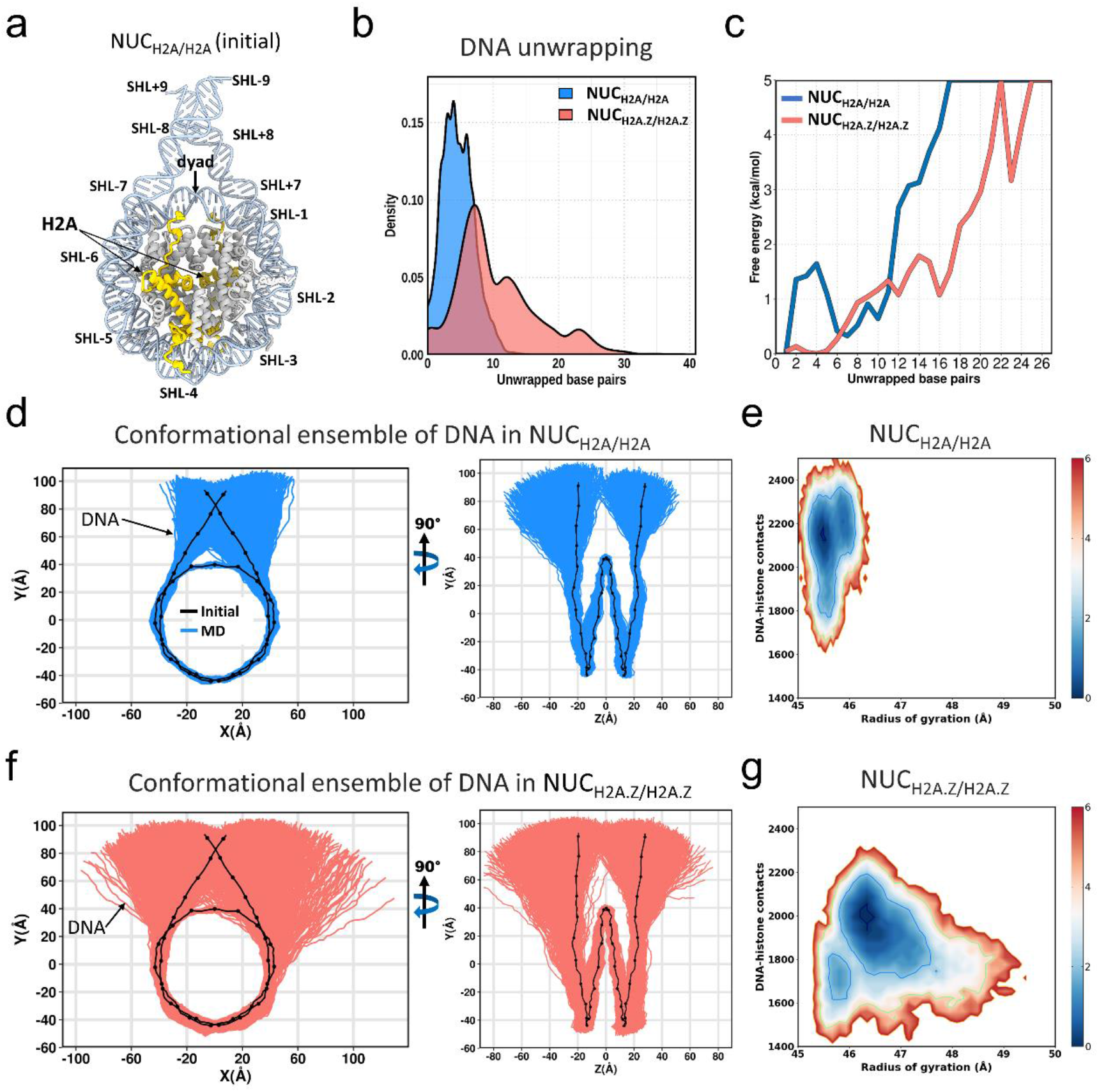
H2A.Z facilitates nucleosomal DNA unwrapping. **a)** Cartoon representation of the initial nucleosome structures for the canonical nucleosome (NUC_H2A/H2A_). **b)** Distributions of a total number of unwrapped base pairs (from an entry DNA side and exit DNA side, see Supplementary Figure S5 for the definitions of entry and exit DNA sides). **c)** Free-energy profile as a function of the unwrapped base pairs for the NUC_H2A/H2A_ (blue) and NUC_H2A.Z/H2A.Z_ (red) nucleosomes. X-axis goes to the maximum number of DNA unwrapped base pairs on one end observed in our simulations (see Supplementary Figure S4 for results for each DNA end from all three simulation runs). **d)** 2D-projections of DNA conformations for NUC_H2A/H2A_ from three independent simulation runs, six microseconds each (see Supplementary Figure S2 for results for each run). **e)** Two-dimensional projection of the free-energy profile as a function of the DNA radius of gyration and the total number of DNA-histone contacts for NUC_H2A/H2A_ nucleosome. **f)** Same as **d)** but for NUC_H2A.Z/H2A.Z_ nucleosome. **g)** Same as **e)** but for NUC_H2A.Z/H2A.Z_ nucleosome. Energy bar values are shown in kcal/mol.

Although the effects of other histone variants are not the focus of our study, we performed six additional MD simulations of two nucleosome systems with the H3.3 variant (Supplementary Figure S1 and Supplementary Table S1). Our results reveal that the homotypic variant nucleosomes with two copies of H2A.Z (NUC_H2A.Z/H2A.Z_ or NUC_H2A.Z/H2A.Z+H3.3/H3.3,_ Figure 1f, Supplementary Figure S3a) exhibit larger magnitudes of DNA unwrapping as compared to heterotypic H2A.Z nucleosomes even with one copy of HA2.Z, no matter how many copies of H3.3 are present (NUC_H2A/H2A.Z+H3/H3.3_, Supplementary Figure S3b). Distributions of the total number of unwrapped base pairs for heterotypic nucleosomes have a characteristic bimodal shape with each peak corresponding to one histone copy (Supplementary Figure S3c). Interestingly, although the distributions of the number of unwrapped base pairs are very similar for NUC_H2A.Z/H2A.Z_ or NUC_H2A.Z/H2A.Z+H3.3/H3.3_ systems, the deposition of H3.3 causes a somewhat more pronounced unwrapping of the entry side of DNA which is less mobile in NUC_H2A.Z/H2A.Z_ systems (Figure 1f). Overall, we can conclude that H2A.Z has the strongest effect in modulating the DNA unwrapping, and the H3.3 variant may play a less prominent role. Our result is in agreement with previous X-ray structure studies showing that incorporation of histone H3.3 did not affect the structure of the H2A.Z-containing nucleosome (23). However, it was also reported that the effect of H2A.Z on nucleosome stability was strongly dependent on the presence of H3.3 (25).

### Energy barrier for DNA unwrapping is reduced in H2A.Z systems

We further computed the free-energy profile as a function of the number of unwrapped base pairs to estimate the energy barriers in the process of DNA unwrapping. Although energy profiles clearly depended on DNA sequence, our results show that in the canonical H2A nucleosome, an energy barrier of ∼1 kcal/mol and ∼5 kcal/mol is needed to unwrap ∼5 and ∼17 base pairs (the maximum degree of unwrapping observed for the canonical system on our time scale) respectively from one end of nucleosomal DNA (Figure 1c, Supplementary Figure S4). This is in good agreement with the DNA unwrapping energy around ∼17 bp reported earlier (33,34). Previous experimental forced disassembly studies on single-molecule unzipping of nucleosomal DNA pointed to the free energy of dissociation of the outside 76 bp being about 12 kcal/mol (35) and found the first energy barrier around the position 17 bp in the canonical nucleosome (13).

Next, we find that the incorporation of variant H2A.Z significantly reduces the energy barrier by several kcal/mol compared to the canonical nucleosome in this range of unwrapping (Figure 1c), consistent with the previous experimental estimates (13). Figures 1e and 1g present the two-dimensional free-energy landscape as a function of the DNA radius of gyration (R_g_) and the total number of DNA-histone contacts. NUC_H2A/H2A_ exhibits a smaller range of R_g_ which is coupled with a larger number of DNA-histone contacts compared to NUC_H2A.Z/H2A.Z_. The landscapes clearly illustrate that the energy cost of opening the two DNA ends in NUC_H2A.Z/H2A.Z_ is considerably smaller than that in NUC_H2A/H2A_. To verify these findings, we performed the MM/GBSA calculations showing that the overall histone-DNA binding energy of the NUC_H2A.Z/H2A.Z_ system is higher than that of the NUC_H2A/H2A_ system (Table S2), indicating the incorporation of H2A.Z component disfavors the DNA-histone binding event, which is consistent with their unwrapping characteristics described above. A much smaller histone-DNA binding energy difference between canonical and H2A.Z nucleosomes was observed previously (36).

### H2A.Z C-terminal tail modulates the DNA unwrapping

In the first section, we showed that DNA ends of H2A.Z-containing nucleosomes are more mobile and are easier to unwrap from the histone octamer when compared to nucleosomes containing canonical H2A. It has been reported earlier that H3 N-terminal and H2A C-terminal tails mediated the unwrapping of DNA ends in nucleosomes (30,31,37-41). Consistent with these findings, we show that the degree of DNA unwrapping is anti-correlated with the number of contacts between the H2A.Z C-terminal tail and the outer DNA region (Figure 2a, see Supplementary Figure S5 for definitions of DNA regions). Therefore, we speculate that the differences in DNA unwrapping between the canonical and H2A.Z nucleosomes might be due to the distinct features in the H2A.Z C-terminal tail. Indeed, in comparison with the H2A C-terminal region (residues 121–130), the H2A.Z C-terminal tail (residues 123–128) lacks the last four residues (including three lysines, Figure 3a) that carry positive charges and may interact with DNA. The analysis of the conformational ensemble of the tails shows that the average distance between the DNA end and C-terminal tail in NUC_H2A/H2A_ is much smaller than that in NUC_H2A.Z/H2A.Z_ (∼3.5 Å vs ∼15 Å, Figure 2c). This indicates that extensive interactions are formed between the H2A C-terminal tail and DNA with contacts predominantly distributed in DNA regions from SHL ± 6 to SHL ± 7 (Figure 2d).

**Figure 2.**
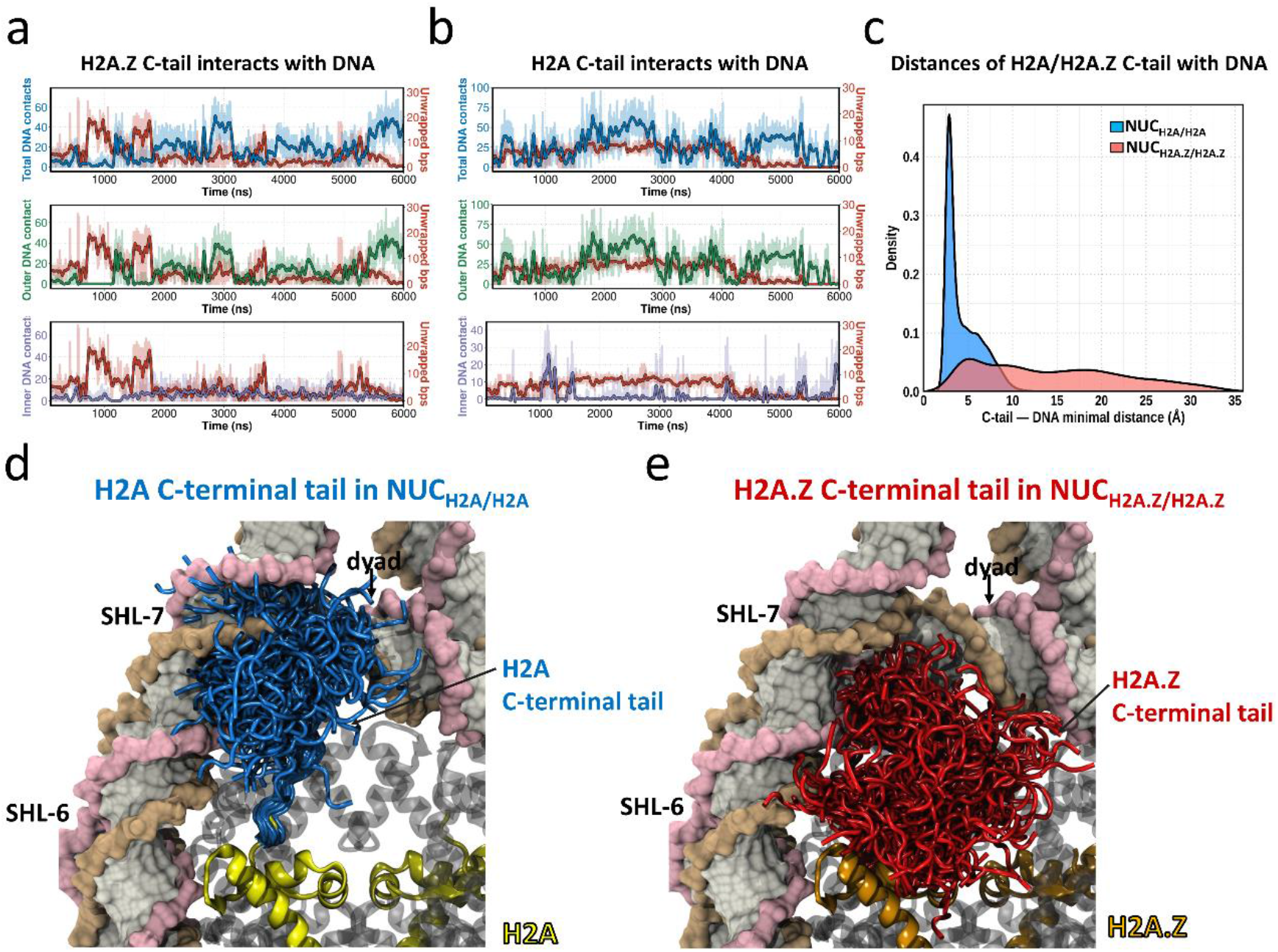
H2A.Z C-terminal tail modulates the DNA unwrapping. **a)** Coupling between the number of unwrapped base pairs from the exit DNA side and the number of H2A.Z C-terminal tail – DNA contacts in NUC_H2A.Z/H2A.Z_ nucleosome. The lines with fade colors are raw data and are smoothed with Savitzky-Golay filter using a ten ns window and first-degree polynomial (dark color lines). **b)** Same as **a)** but for NUC_H2A/H2A_ nucleosome. **c)** The distribution of distances between the DNA end segment (bps ±41 to ±75) and H2A or H2A.Z C-terminal tail (H2A: residues 1 to16; H2A.Z: residues 1 to18) from three independent simulation runs. **d)** Structural conformations of H2A C-terminal tail (blue color, residues 121–130) in canonical nucleosome (NUC_H2A/H2A_) from three simulation runs. **e)** Structural conformations of H2A.Z C-terminal tail (red color, residues 123–128) in H2A.Z nucleosome (NUC_H2A.Z/H2A.Z_) from three simulation runs. All frames are taken based on 50ns interval.

**Figure 3.**
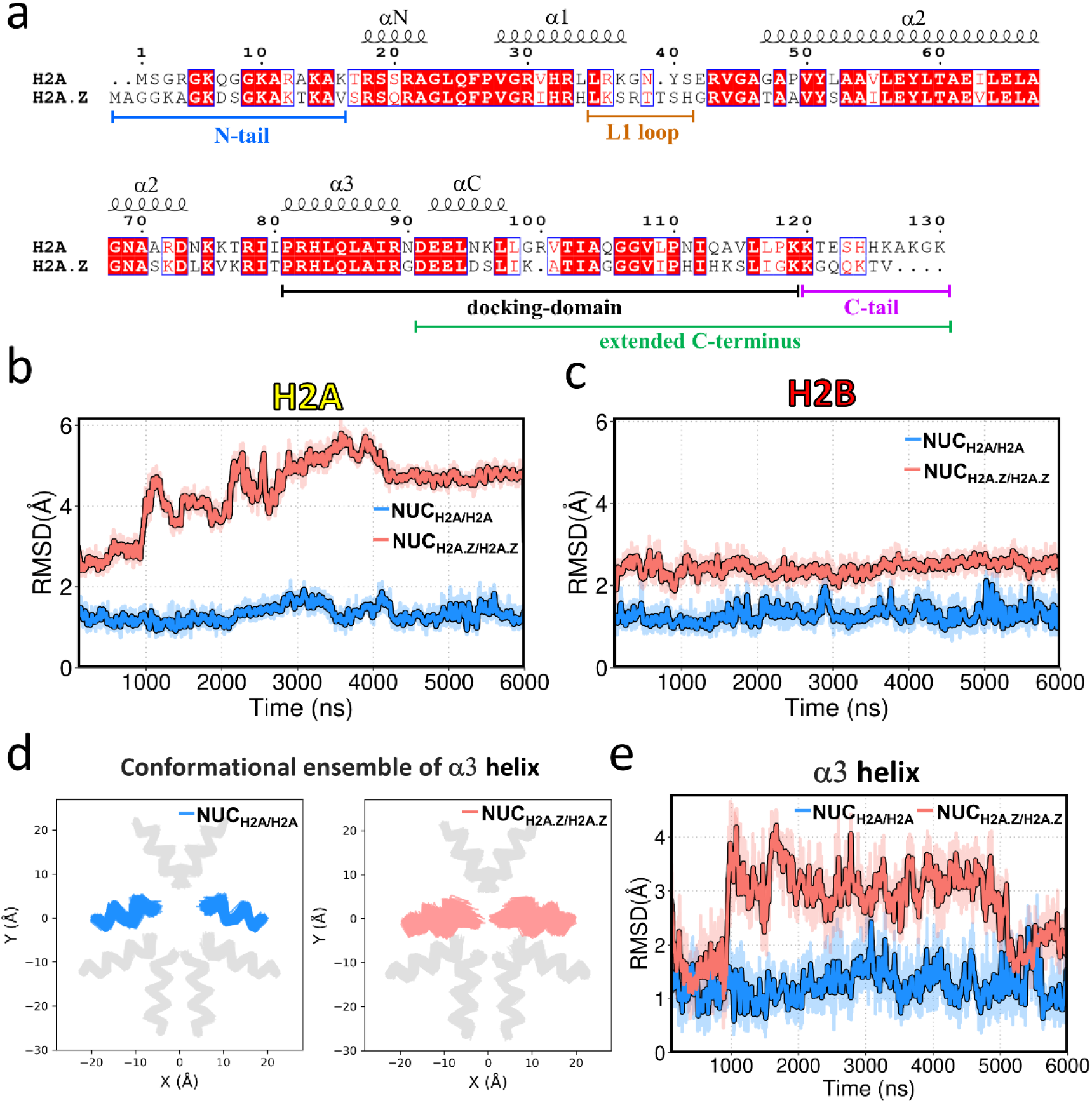
H2A.Z incorporation increases the plasticity of the histone octamer. **a)** Sequence comparison between H2A and H2A.Z. **b)** RMSD of Cα-atoms as a function of simulation time for histone H2A core region in NUC_H2A/H2A_ nucleosome (blue) and H2A.Z core region in NUC_H2A.Z/H2A.Z_ nucleosome (red). The lines with fade colors are raw data and are smoothed with Savitzky-Golay filter using a ten ns window and first-degree polynomial (dark color lines). **c)** RMSD of Cα-atoms as a function of simulation time for histone H2B core regions in NUC_H2A/H2A_ nucleosome (blue) and NUC_H2A.Z/H2A.Z_ nucleosome (red). **d)** Conformational ensemble of H2A α3 helix in NUC_H2A/H2A_ nucleosome (blue) and H2A.Z α3 helix in NUC_H2A.Z/H2A.Z_ nucleosome (red). Other types of histones are shown in grey. See Supplementary Figure S9 for results of other helices. **e)** RMSD of Cα-atoms as a function of simulation time for H2A α3 helix in NUC_H2A/H2A_ nucleosome (blue) and H2A.Z α3 helix in NUC_H2A.Z/H2A.Z_ nucleosome (red).

In contrast, the distance distribution between the H2A.Z C-terminal tail and DNA end is substantially shifted to larger values and can reach 35 Å (Figure 2c). As the result of the loss of interactions with DNA, the H2A.Z C-terminal tail almost entirely protrudes from the nucleosome disc into the solvent with only a few tail conformations interacting with the DNA end (Figure 2f). These results suggest that the absence of positively charged residues at the end of the tail in H2A.Z dramatically weakens the association between the H2A.Z C-terminal tail and DNA, leading to the unwrapping of DNA. Consistent with this result, a recent single-particle cryo-EM study suggested that the H2A.Z C-terminal tail was more flexible than the canonical H2A (42). Interestingly, we find that the protruded C-terminal tail of H2A.Z is long enough to interact with an adjacent nucleosome (Supplementary Figure S6). Namely, a representative H2A.Z C-terminal tail conformation (with the extended C-terminus, see Figure 3a) can extend to distances up to 29 Å, which is much larger than the nucleosome-nucleosome distance in the nucleosome arrays (43).

### H2A.Z deposition increases the plasticity of the histone octamer

The structural dynamics of the histone octamer backbone with the incorporation of H2A.Z may be also associated with DNA unwrapping. Figures 3b and 3c show the time evolution of the displacement of different regions of core H2A/H2A.Z and H2B (without histone tails) with respect to the equilibrated structure, measured as RMSD. Although there is no obvious difference in RMSD of Cα-atoms for histone H3 and H4 core regions between NUC_H2A/H2A_ and NUC_H2A.Z/H2A.Z_ nucleosomes (Supplementary Figure S7), a noticeable increase in dynamics is observed for H2A.Z core regions in NUC_H2A.Z/H2A.Z_ (up to 6 Å) compared to the canonical H2A in NUC_H2A/H2A_ (Figure 3b). Interestingly, RMSD of histone H2B is also higher in NUC_H2A.Z/H2A.Z_ nucleosomes (Figure 3c) possibly because of the dimer formation between H2A.Z and H2B and long-range distance effects. Moreover, the α3 helix of H2A.Z which is located within the docking domain and near the C-terminus, exhibits relatively larger RMSD values than α1 and α2 helices (Figure 3e, Supplementary Figure S8). We further plotted the 2D projections of individual H2A and H2A.Z α-helices and find that the increased dynamics of H2A.Z core regions in NUC_H2A.Z/H2A.Z_ are characterized by much larger fluctuations (Figure 3d, Supplementary Figure S9). This result suggests that the overall octamer plasticity is enhanced in the H2A.Z nucleosome, which in turn can be associated with the nucleosomal and linker DNA dynamics. Several recent studies revealed that restraining histone octamer plasticity indeed suppressed DNA fluctuations and led to a lower magnitude of DNA unwrapping and breathing (31,44,45).

### Gaping motions are enhanced in H2A.Z nucleosomes

In addition to the enhanced DNA unwrapping induced by the incorporation of histone variant H2A.Z, another striking alteration is the noticeable spontaneous nucleosome gaping. Nucleosome gaping motions are distinct from DNA unwrapping. During nucleosome gaping transitions, two half-turns (SHL+ and SHL-) of nucleosomal DNA open apart by 5 -10 Å along the axis which is perpendicular to the nucleosome plane (46). To quantitatively characterize the nucleosome gapping, we measured the distance between the two DNA segments in SHL-4 and SHL+4 regions (Figure 4a). In the canonical nucleosome, no significant gaping event is observed and the distances do not exceed ∼24 Å. However, the distance distribution between two DNA gyres is shifted to larger values in the H2A.Z nucleosomes with two peaks at ∼ 27 Å and 36 Å, suggesting the nucleosome gaping motions are substantially enhanced by the H2A.Z variant. Previous studies showed that during the DNA translocation of chromatin remodeler SWR1, the gaping distances measured between SHL±2 widened roughly from 27 Å to 36 Å in the presence of SWR1 (47). Although these studies were conducted on canonical nucleosomes, the spontaneous gapping could be even more important for binding chromatin remodellers by H2A.Z containing nucleosomes.

**Figure 4.**
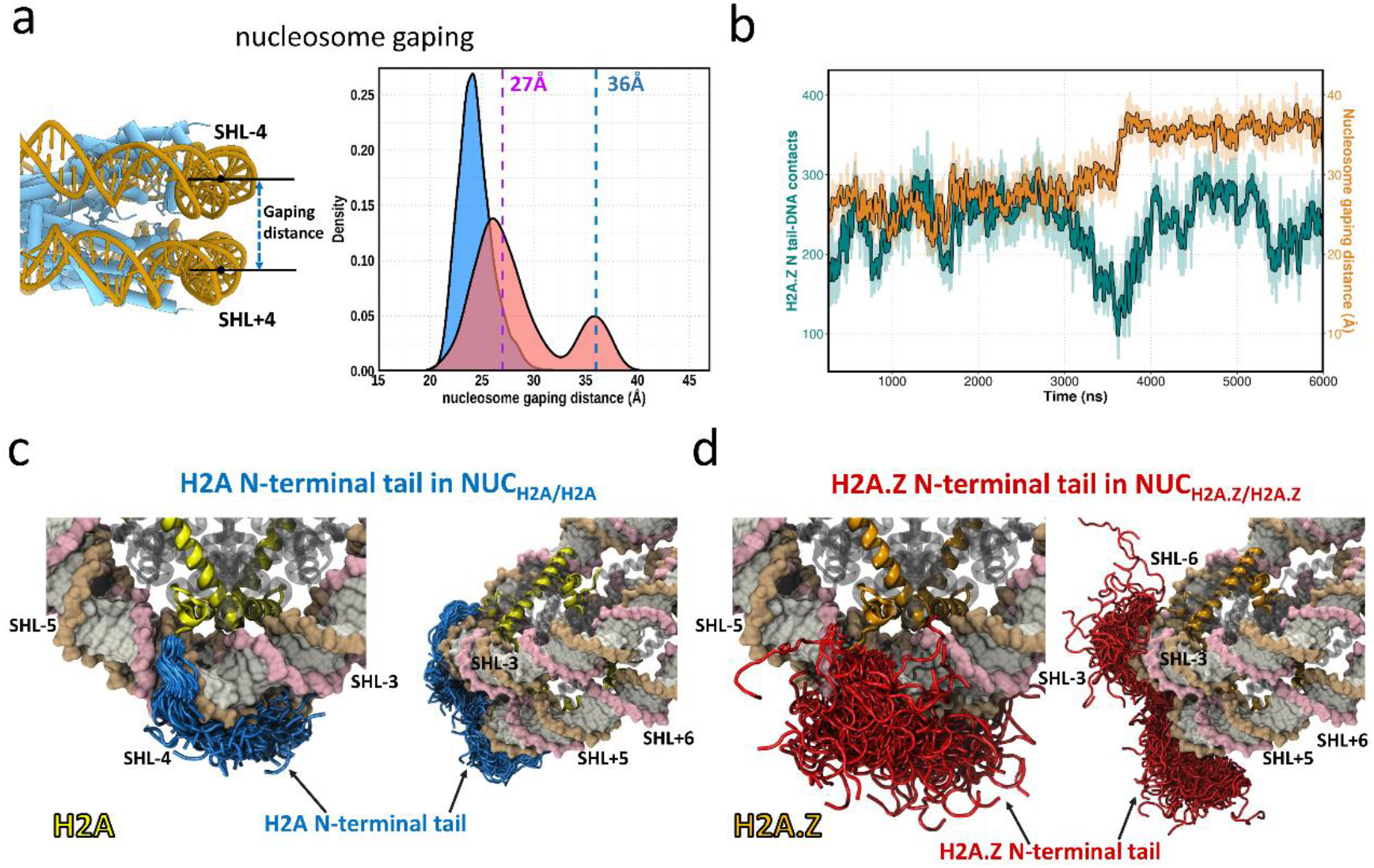
H2A.Z N-terminal tail modulates nucleosome gapping. **a)** Distributions of nucleosome gaping distances in NUC_H2A/H2A_ (blue) and NUC_H2A.Z/H2A.Z_ (red) systems from three independent simulation runs. Vertical dashed lines indicate the gaping distances at SHL±2 in the presence of SWR1 during the DNA translocation (47). **b)** Time evolution of the H2A.Z N-terminal tail - DNA contacts (green) and nucleosome gapping distances (orange) in NUC_H2A.Z/H2A.Z_. See Supplementary Figure S10 for the H2A N-terminal tail - DNA contacts and nucleosome gapping distances in NUC_H2A/H2A_ system. **c)** Structural conformations of H2A N-terminal tail (blue color, residues 1-16) in the canonical nucleosome (NUC_H2A/H2A_) from three simulation runs. **d)** Structural conformations of H2A.Z N-terminal tail (red color, residues 1-18) in H2A.Z nucleosome (NUC_H2A.Z/H2A.Z_) from three simulation runs. All frames are taken based on 50ns interval.

To explore the molecular mechanism of H2A.Z-induced nucleosome gapping, we calculated the time evolution of the number of contacts between the H2A.Z N-terminal tail (residues 1-18) and DNA and examined its correlation with the gaping distances (Figure 4b). We find that the notable nucleosome gapping events are tightly coupled with the decreased number of contacts between the H2A.Z N-terminal tail and DNA. Namely, a striking decrease in the number of contacts is observed at around 3.5 µs, followed by a significant nucleosome gapping transition into the second state with a 36 Å distance between the two gyres. This state is stable enough to last for another several microseconds despite the reestablishment of the contacts. This result indicates that the enhanced nucleosome gapping in the H2A.Z nucleosome may be caused by the loss of interactions between DNA and the H2A.Z N-terminal tail. From the conformational analysis of N-terminal tails, we observe that N-terminal tails of H2A are predominantly inserted into the DNA minor and major grooves between SHL±3 and SHL±4 (Figure 4c), which slightly shifted from their original binding mode in H2A nucleosomes (37). In contrast, the H2A.Z N-terminal tails have a preference to be exposed to the solvent and exhibit a relatively large conformational variability and have a reduced number of contacts with DNA (Figure 4d). It can be explained by the difference in their net charge, with the H2A N-terminal tail carrying a higher positive charge (by 2+) compared to the H2A.Z tail. Interestingly, a recent study reported that the N-terminal region of H2A.Z.1 (residue 1-60) is responsible for the flexible H2A.Z nucleosome positioning, possibly because of the reduced histone–DNA interactions (48).

## Discussion

Histone variants play crucial roles in the evolution of complex multicellular organisms and expand the function of chromatin. (42,49). It has been debated for a long time whether nucleosomes containing the H2A.Z variant are thermodynamically and dynamically stable or unstable because shreds of evidence exist in support of stabilization (50) or destabilization (23,42,51). In the present study, we estimated the free energy barriers and performed multiple runs of six-microsecond all-atom MD simulations to characterize the detailed atomistic mechanism by which human H2A.Z influences the dynamics and stability of nucleosomes. We find that the incorporation of the H2A.Z variant facilitates the DNA unwrapping from the octamer and nucleosome gaping and increases the histone octamer dynamics. Importantly, this is the first time that spontaneous DNA unwrapping for up to forty base pairs on both ends is observed for full nucleosomes with histone tails in an all-atom MD simulation. Spontaneous DNA unwrapping in canonical nucleosomes on a time scale of tens and hundreds of milliseconds was recorded experimentally under physiological solution conditions (52-55). The difference in time scales of these motions observed in our simulations and previous experimental studies (mostly FRET) can be explained by the higher mobility of H2A.Z nucleosomes compared to the canonical ones, by the native DNA sequence used in our study, rather than the well-positioned 601 sequence, and by overlooking the short-lived unwrapped states by FRET, biasing results to a longer lifetime, as reported before (53).

It has been known for a while that certain histone tails form interactions with the nucleosomal and linker DNA and suppress DNA unwrapping (37,42,56-59). Variant histones carry many alterations in tails including amino acid substitutions, insertions or deletions. Perhaps, the deposition of H2A.Z is coupled with its destabilization role in enhancing nucleosome dynamics and DNA unwrapping through the means of histone tails. Indeed, our results show that a shorter and less positively charged C-terminal tail of H2A.Z leads to a significant interaction disruption with the DNA ends as compared to canonical nucleosomes, which contributes to DNA unwrapping. Moreover, there is a possibility that in the nucleosomal array context, the freed H2A.Z C-terminal tails may form interactions with the increased acidic patch from the neighboring H2A.Z nucleosome or mediate fiber-fiber interactions as suggested previously (60), enabling the formation of more compact chromatin fiber compared to the canonical systems, which can potentially reconcile the role of H2A.Z in both transcription activation and facilitation of the intramolecular folding of nucleosomal arrays (61). On the other hand, the H2A.Z N-terminal tail might be responsible for the enhanced nucleosome gaping transition, because of weakened DNA-histone interactions.

It is reasonable to speculate that the connection between the increased mobility of H2A.Z nucleosomes and their function is to make nucleosomes more mobile and DNA more accessible to transcriptional machinery and other chromatin components. It has been known that histone mutations (mostly involving H3 and H2A) that alter the DNA entry-exit site accessibility have also direct effects on transcription elongation and termination (62). Indeed, it has been suggested previously that H2A.Z incorporation enhances terminal DNA accessibility (42) and facilitates transcription initiation (63,64). This is also consistent with earlier studies showing that H2A.Z nucleosomes protect only 120 base pairs of nucleosomal DNA (65).

Taken together, our main results may provide a detailed explanation to support the crucial role of H2A.Z in transcription following the model suggested previously (11,12,66,67). According to the latter, the promoter-distal half of the +1 nucleosome presents a strong barrier to transcription (66,67), and the H2A.Z variant is enriched on this site and reduces the nucleosomal barrier at the transcription start site (11,12). According to our estimates, the energy barrier is reduced by ∼4 kcal/mol in H2A.Z compared to canonical nucleosomes. The major role in this mechanism is played by H2A.Z tails, which lose many contacts with DNA. This enables the asymmetric unwrapping and gaping of DNA in H2A.Z containing nucleosomes. Such increased dynamics, in turn, can guide the octasome-to-hexasome transitions (68). Overall, our study provides insights into dissecting the major structural and dynamic factors that play important roles in H2A.Z containing nucleosomes. However, one has to keep in mind that their function depends on many other factors, including the presence of histone and DNA covalent modifications and genomic location.

## Methods

### Nucleosome modeling with the human native DNA sequence

The simulated systems in this study consist of four full nucleosome models with two straight 20-bp long DNA linker segments. The 187 bp long (147+2×20) DNA sequence is taken from *Homo sapiens TP53* gene +1 nucleosome. The X-ray structure of the human nucleosome core particle (PDB ID: 3AFA) was used to construct the canonical homotypic nucleosome containing two copies of H2A. The detailed procedure for constructing the nucleosome models with the native DNA sequence can be found in our recently published paper (37). The human homotypic H2A.Z nucleosome model was built based on the X-ray structure (PDB ID: 1F66) containing human H2A.Z.1, histones H3, H4, and H2B were substituted by canonical human histone sequences. Considering that histone variant H3.3 and heterotypic histone dimers such as H2A/H2A.Z also play essential functional and structural roles (25), we also constructed structural models to include H3.3 or H2A/H2A.Z based on two other X-ray crystal structures (PDB IDs: 5B33 and 5B32). The histone tails were linearly extended into the solvent symmetrically oriented with respect to the dyad axis. Thus, totally four nucleosome models were constructed: NUC_H2A/H2A_ (H2A/H2A+H3/H3), NUC_H2A.Z/H2A.Z_ (H2A.Z/H2A.Z+H3/H3), NUC_H2A.Z/H2A.Z+H3.3/H3.3_(H2A.Z/H2A.Z+H3.3/H3.3), and NUC_H2A/H2A.Z+H3/H3.3_ (H2A/H2A.Z+H3/H3.3). For each model, three independent simulation runs were performed with each run trajectory extending up to six microseconds totaling 72 microseconds of simulation time.

### Molecular dynamics simulation protocol

All MD simulations were performed with the package GROMACS version 2020.6 (69) using the AMBER ff14SB force field for protein and OL15 for double-stranded DNA (70,71). Simulations were conducted in explicit solvent using an Optimal Point Charge (OPC) water model, which was recently illustrated to reproduce water liquid bulk properties and to greatly improve nucleic acid simulations as well as intrinsically disordered protein simulations. (72). The OPC water model was used in our previous long simulations of histone tails and kinetic and thermodynamic estimates from these simulations were shown to be reasonable and agreed with observables (73). In each simulation system, the initial structural model was solvated in a box with a minimum distance of 20 Å between the nucleosome atoms and the edges of the box. NaCl was added to the system up to a concentration of 150 mM. The solvated systems were first energy minimized using steepest descent minimization for 10,000 steps, gradually heated to 310 K throughout 800 ps using restraints, and then equilibrated for a period of 1 ns. After that, the production simulations were carried out in the isobaric-isothermic (NPT) ensemble up to 6 µs, with the temperature maintained at 310 K using the modified Berendsen thermostat (velocity-rescaling) (74) and the pressure maintained at 1 atm using the Parrinello–Rahman barostat (75). A cutoff of 10 Å was applied to short-range non-bonded vdW interactions, and the Particle Mesh Ewald (PME) method (76) was used to calculate all long-range electrostatic interactions. Periodic boundary conditions were used. Covalent bonds involving hydrogens were constrained to their equilibrium lengths using the LINCS algorithm (77), allowing a 2 fs time step to be employed. Coordinates of the solutes were collected every 100 ps yielding a total of 60,000 frames for further analysis. The set of all simulated systems and their parameters are provided in Supplementary Table 1.

### Analysis of nucleosome dynamics

MD trajectory snapshots were first processed by performing a root mean square deviation (RMSD) fit of the C-α atoms of the histone core to the minimized structure of the nucleosome. The first 200 nanosecond frames of each simulation were treated as equilibration periods and were excluded from the analysis. In-house codes were developed to quantify the DNA and histone paths, and DNA-histone interactions. The interactions were defined by two non-hydrogen atoms from DNA and histone within a distance of less than 4.5 Å. The unwrapped base pairs were defined as those in which the center of the base pair deviates more than 7 Å from the corresponding base pair in the initial structure. The radius of gyration of DNA segments was calculated using gmx_gyrate (69). The free energy of DNA unwrapping was calculated using the relation ΔG = −RT ln (f/f_max_), where f is the frequency of a particular number of unwrapped base pairs and f_max_ is the maximum frequency found for the unwrapped base pairs. For example, the frequency of unwrapped base pairs at 10 f_10_=N_10_/N_total_, where N_10_ is the total number of frames with 10 unwrapped base pairs in the MD simulation trajectory, N_total_ is the total number of frames in the trajectory (60000). The multi-dimensional free energy landscapes were calculated using gmx_sham (69) with the radius of gyration and the number of histone-DNA contacts as collective variables. The binding free energy between the histone octamer and the native DNA sequence was also calculated using the molecular mechanics generalized Born surface area (MM/GBSA) method, which is embedded in the Amber20 Package (78). The MM/GBSA free energy calculation on both NUC_H2A/H2A_ and NUC_H2A.Z/H2A.Z_ systems was based on the first 200ns production trajectories. A residue-wise free energy decomposition was conducted to access the energy profile of both DNA ends, where the interaction energy of every DNA residue with every other atom in the system was calculated. Note that the MM/GBSA approach tends to overestimate the absolute energy. In a previous paper, Peng et al. showed that there is high correlation coefficients between the ΔG_0_ estimated from MM/GBSA calculations and the conformational ensemble statistics (37). However, both methods provide a semi-qualitative estimation of the free energy profile of DNA unwrapping.

## Acknowledgments

We would like to thank Tiina Liinamaa for proofreading the manuscript. SL, TW and ARP were supported by the Department of Pathology and Molecular Medicine, Queen’s University, Canada. ARP is the recipient of a Senior Canada Research Chair in Computational Biology and Biophysics and a Senior Investigator Award from the Ontario Institute of Cancer Research, Canada. ARP and SL acknowledge the support of the Natural Sciences and Engineering Research Council of Canada (NSERC). This study used high-performance computational resources from Compute Canada (https://docs.computecanada.ca).

## References

1. Luger, K., Mäder, A.W., Richmond, R.K., Sargent, D.F. and Richmond, T.J. (1997) Crystal structure of the nucleosome core particle at 2.8 Å resolution. Nature, 389, 251–260.

2. Talbert, P.B., Ahmad, K., Almouzni, G., Ausió, J., Berger, F., Bhalla, P.L., Bonner, W.M., Cande, W.Z., Chadwick, B.P., Chan, S.W.L. et al. (2012) A unified phylogeny-based nomenclature for histone variants. Epigenetics & Chromatin, 5, 7.

3. Draizen, E.J., Shaytan, A.K., Marino-Ramirez, L., Talbert, P.B., Landsman, D. and Panchenko, A.R. (2016) HistoneDB 2.0: a histone database with variants-an integrated resource to explore histones and their variants. Database-Oxford.

4. Raisner, R.M., Hartley, P.D., Meneghini, M.D., Bao, M.Z., Liu, C.L., Schreiber, S.L., Rando, O.J. and Madhani, H.D. (2005) Histone Variant H2A.Z Marks the 5′ Ends of Both Active and Inactive Genes in Euchromatin. Cell, 123, 233–248.

5. Zhang, H., Roberts, D.N. and Cairns, B.R. (2005) Genome-Wide Dynamics of Htz1, a Histone H2A Variant that Poises Repressed/Basal Promoters for Activation through Histone Loss. Cell, 123, 219–231.

6. Albert, I., Mavrich, T.N., Tomsho, L.P., Qi, J., Zanton, S.J., Schuster, S.C. and Pugh, B.F. (2007) Translational and rotational settings of H2A.Z nucleosomes across the Saccharomyces cerevisiae genome. Nature, 446, 572–576.

7. Giaimo, B.D., Ferrante, F., Herchenröther, A., Hake, S.B. and Borggrefe, T. (2019) The histone variant H2A.Z in gene regulation. Epigenetics & Chromatin, 12, 37.

8. Ibarra-Morales, D., Rauer, M., Quarato, P., Rabbani, L., Zenk, F., Schulte-Sasse, M., Cardamone, F., Gomez-Auli, A., Cecere, G. and Iovino, N. (2021) Histone variant H2A.Z regulates zygotic genome activation. Nature Communications, 12.

9. Moosmann, A., Campsteijn, C., Jansen, P.W.T.C., Nasrallah, C., Raasholm, M., Stunnenberg, H.G. and Thompson, E.M. (2011) Histone variant innovation in a rapidly evolving chordate lineage. Bmc Evol Biol, 11.

10. Gursoy-Yuzugullu, O., Ayrapetov, M.K. and Price, B.D. (2015) Histone chaperone Anp32e removes H2A.Z from DNA double-strand breaks and promotes nucleosome reorganization and DNA repair. P Natl Acad Sci USA, 112, 7507–7512.

11. Weber, Christopher M., Ramachandran, S. and Henikoff, S. (2014) Nucleosomes Are Context-Specific, H2A.Z-Modulated Barriers to RNA Polymerase. Molecular Cell, 53, 819–830.

12. Rudnizky, S., Bavly, A., Malik, O., Pnueli, L., Melamed, P. and Kaplan, A. (2016) H2A.Z controls the stability and mobility of nucleosomes to regulate expression of the LH genes. Nature Communications, 7.

13. Chen, Z.J., Gabizon, R., Brown, A.I., Lee, A., Song, A.X., Diaz-Celis, C., Kaplan, C.D., Koslover, E.F., Yao, T.T. and Bustamante, C. (2019) High-resolution and high-accuracy topographic and transcriptional maps of the nucleosome barrier. Elife, 8.

14. Mylonas, C., Lee, C., Auld, A.L., Cisse, I.I. and Boyer, L.A. (2021) A dual role for H2AZ1 in modulating the dynamics of RNA polymerase II initiation and elongation. Nature Structural & Molecular Biology, 28, 435-+.

15. Hardy, S., Jacques, P.-É., Gévry, N., Forest, A., Fortin, M.-È., Laflamme, L., Gaudreau, L. and Robert, F. (2009) The Euchromatic and Heterochromatic Landscapes Are Shaped by Antagonizing Effects of Transcription on H2A.Z Deposition. Plos Genet, 5, e1000687.

16. Lashgari, A., Millau, J.-F., Jacques, P.-É. and Gaudreau, L. (2017) Global inhibition of transcription causes an increase in histone H2A.Z incorporation within gene bodies. Nucleic Acids Research, 45, 12715–12722.

17. Babiarz, J.E., Halley, J.E. and Rine, J. (2006) Telomeric heterochromatin boundaries require NuA4-dependent acetylation of histone variant H2A.Z in Saccharomyces cerevisiae. Gene Dev, 20, 700–710.

18. Rangasamy, D., Greaves, I. and Tremethick, D.J. (2004) RNA interference demonstrates a novel role for H2A.Z in chromosome segregation. Nature Structural & Molecular Biology, 11, 650–655.

19. Xu, Y., Ayrapetov, M.K., Xu, C., Gursoy-Yuzugullu, O., Hu, Y.D. and Price, B.D. (2012) Histone H2A.Z Controls a Critical Chromatin Remodeling Step Required for DNA Double-Strand Break Repair. Molecular Cell, 48, 723–733.

20. Sales-Gil, R., Kommer, D.C., de Castro, I.J., Amin, H.A., Vinciotti, V., Sisu, C. and Vagnarelli, P. (2021) Non-redundant functions of H2AZ1 and H2AZ2 in chromosome segregation and cell cycle progression. EMBO Rep, 22.

21. Abbott, D.W., Ivanova, V.S., Wang, X., Bonner, W.M. and Ausió, J. (2001) Characterization of the Stability and Folding of H2A.Z Chromatin Particles: IMPLICATIONS FOR TRANSCRIPTIONAL ACTIVATION *. Journal of Biological Chemistry, 276, 41945–41949.

22. Watanabe, S., Radman-Livaja, M., Rando Oliver, J. and Peterson Craig, L. (2013) A Histone Acetylation Switch Regulates H2A.Z Deposition by the SWR-C Remodeling Enzyme. Science, 340, 195–199.

23. Horikoshi, N., Arimura, Y., Taguchi, H. and Kurumizaka, H. (2016) Crystal structures of heterotypic nucleosomes containing histones H2A.Z and H2A. Open Biol, 6.

24. Suto, R.K., Clarkson, M.J., Tremethick, D.J. and Luger, K. (2000) Crystal structure of a nucleosome core particle containing the variant histone H2A.Z. Nature Structural Biology, 7, 1121–1124.

25. Jin, C. and Felsenfeld, G. (2007) Nucleosome stability mediated by histone variants H3.3 and H2A.Z. Genes Dev, 21, 1519–1529.

26. Wen, Z.Q., Zhang, L.W., Ruan, H.H. and Li, G.H. (2020) Histone variant H2A.Z regulates nucleosome unwrapping and CTCF binding in mouse ES cells. Nucleic Acids Research, 48, 5939–5952.

27. Draker, R., Ng, M.K., Sarcinella, E., Ignatchenko, V., Kislinger, T. and Cheung, P. (2012) A Combination of H2A.Z and H4 Acetylation Recruits Brd2 to Chromatin during Transcriptional Activation. Plos Genet, 8.

28. Jin, C.Y., Zang, C.Z., Wei, G., Cui, K.R., Peng, W.Q., Zhao, K.J. and Felsenfeld, G. (2009) H3.3/H2A.Z double variant-containing nucleosomes mark ‘nucleosome-free regions’ of active promoters and other regulatory regions. Nat Genet, 41, 941–U112.

29. Ngo, T.T., Zhang, Q., Zhou, R., Yodh, J.G. and Ha, T. (2015) Asymmetric unwrapping of nucleosomes under tension directed by DNA local flexibility. Cell, 160, 1135–1144.

30. Li, S.X., Peng, Y.H., Landsman, D. and Panchenko, A.R. (2022) DNA methylation cues in nucleosome geometry, stability and unwrapping. Nucleic Acids Research, 50, 1864–1874.

31. Armeev, G.A., Kniazeva, A.S., Komarova, G.A., Kirpichnikov, M.P. and Shaytan, A.K. (2021) Histone dynamics mediate DNA unwrapping and sliding in nucleosomes. Nat Commun, 12, 2387.

32. Winogradofft, D. and Aksimentiev, A. (2019) Molecular Mechanism of Spontaneous Nucleosome Unraveling. Journal of Molecular Biology, 431, 323–335.

33. Zhang, B., Zheng, W.H., Papoian, G.A. and Wolynes, P.G. (2016) Exploring the Free Energy Landscape of Nucleosomes. Journal of the American Chemical Society, 138, 8126–8133.

34. Kono, H., Sakuraba, S. and Ishida, H. (2018) Free energy profiles for unwrapping the outer superhelical turn of nucleosomal DNA. Plos Computational Biology, 14.

35. Brower-Toland, B.D., Smith, C.L., Yeh, R.C., Lis, J.T., Peterson, C.L. and Wang, M.D. (2002) Mechanical disruption of individual nucleosomes reveals a reversible multistage release of DNA. P Natl Acad Sci USA, 99, 1960–1965.

36. Bowerman, S. and Wereszczynski, J. (2016) Effects of MacroH2A and H2A.Z on Nucleosome Dynamics as Elucidated by Molecular Dynamics Simulations. Biophysical Journal, 110, 327–337.

37. Peng, Y.H., Li, S.X., Onufriev, A., Landsman, D. and Panchenko, A.R. (2021) Binding of regulatory proteins to nucleosomes is modulated by dynamic histone tails. Nat Commun, 12, 5280.

38. Huertas, J., Scholer, H.R. and Cojocaru, V. (2021) Histone tails cooperate to control the breathing of genomic nucleosomes. Plos Comput Biol, 17, e1009013.

39. Shaytan, A.K., Armeev, G.A., Goncearenco, A., Zhurkin, V.B., Landsman, D. and Panchenko, A.R. (2016) Coupling between Histone Conformations and DNA Geometry in Nucleosomes on a Microsecond Timescale: Atomistic Insights into Nucleosome Functions. J Mol Biol, 428, 221–237.

40. Li, Z.H. and Kono, H. (2016) Distinct Roles of Histone H3 and H2A Tails in Nucleosome Stability. Sci Rep-Uk, 6.

41. Chakraborty, K. and Loverde, S.M. (2017) Asymmetric breathing motions of nucleosomal DNA and the role of histone tails. J Chem Phys, 147.

42. Lewis, T.S., Sokolova, V., Jung, H., Ng, H. and Tan, D.Y. (2021) Structural basis of chromatin regulation by histone variant H2A.Z. Nucleic Acids Research, 49, 11379–11391.

43. Schalch, T., Duda, S., Sargent, D.F. and Richmond, T.J. (2005) X-ray structure of a tetranucleosome and its implications for the chromatin fibre. Nature, 436, 138–141.

44. Bilokapic, S., Strauss, M. and Halic, M. (2018) Histone octamer rearranges to adapt to DNA unwrapping. Nature Structural & Molecular Biology, 25, 101-+.

45. Hada, A., Hota, S.K., Luo, J., Lin, Y.-c., Kale, S., Shaytan, A.K., Bhardwaj, S.K., Persinger, J., Ranish, J., Panchenko, A.R. et al. (2019) Histone Octamer Structure Is Altered Early in ISW2 ATP-Dependent Nucleosome Remodeling. Cell Reports, 28, 282-294.e286.

46. Ngo, T.T.M. and Ha, T. (2015) Nucleosomes undergo slow spontaneous gaping. Nucleic Acids Research, 43, 3964–3971.

47. Willhoft, O., Ghoneim, M., Lin, C.-L., Chua, E.Y.D., Wilkinson, M., Chaban, Y., Ayala, R., McCormack, E.A., Ocloo, L., Rueda, D.S. et al. (2018) Structure and dynamics of the yeast SWR1-nucleosome complex. Science, 362, eaat7716.

48. Sato, S., Tanaka, N., Arimura, Y., Kujirai, T. and Kurumizaka, H. (2020) The N-terminal and C-terminal halves of histone H2A.Z independently function in nucleosome positioning and stability. Genes Cells, 25, 538–546.

49. Horikoshi, N., Kujirai, T., Sato, K., Kimura, H. and Kurumizaka, H. (2019) Structure-based design of an H2AZ1 mutant stabilizing a nucleosome in vitro and in vivo. Biochem Bioph Res Co, 515, 719–724.

50. Hoch, D.A., Stratton, J.J. and Gloss, L.M. (2007) Protein-protein forster resonance energy transfer analysis of nucleosome core particles containing H2A and H2A.Z. Journal of Molecular Biology, 371, 971–988.

51. Bönisch, C., Schneider, K., Pünzeler, S., Wiedemann, S.M., Bielmeier, C., Bocola, M., Eberl, H.C., Kuegel, W., Neumann, J., Kremmer, E. et al. (2012) H2A.Z.2.2 is an alternatively spliced histone H2A.Z variant that causes severe nucleosome destabilization. Nucleic Acids Research, 40, 5951–5964.

52. Li, G., Levitus, M., Bustamante, C. and Widom, J. (2005) Rapid spontaneous accessibility of nucleosomal DNA. Nature Structural & Molecular Biology, 12, 46–53.

53. Koopmans, W.J.A., Brehm, A., Logie, C., Schmidt, T. and van Noort, J. (2007) Single-pair FRET microscopy reveals mononucleosome dynamics. J Fluoresc, 17, 785–795.

54. Poirier, M.G., Bussiek, M., Langowski, J. and Widom, J. (2008) Spontaneous access to DNA target sites in folded chromatin fibers. Journal of Molecular Biology, 379, 772–786.

55. Poyton, M.F., Feng, X.A., Ranjan, A., Lei, Q., Wang, F., Zarb, J.S., Louder, R.K., Park, G., Jo, M.H., Ye, J. et al. (2022) Coordinated DNA and histone dynamics drive accurate histone H2A.Z exchange. Sci Adv, 8.

56. Brower-Toland, B., Wacker, D.A., Fulbright, R.M., Lis, J.T., Kraus, W.L. and Wang, M.D. (2005) Specific contributions of histone tails and their acetylation to the mechanical stability of nucleosomes. Journal of Molecular Biology, 346, 135–146.

57. Peng, Y.H., Li, S.X., Landsman, D. and Panchenko, A.R. (2021) Histone tails as signaling antennas of chromatin. Current Opinion in Structural Biology, 67, 153–160.

58. Ali-Ahmad, A., Bilokapic, S., Schafer, I.B., Halic, M. and Sekulic, N. (2019) CENP-C unwraps the human CENP-A nucleosome through the H2A C-terminal tail. EMBO Rep, 20.

59. Vogler, C., Huber, C., Waldmann, T., Ettig, R., Braun, L., Izzo, A., Daujat, S., Chassignet, I., Lopez-Contreras, A.J., Fernandez-Capetillo, O. et al. (2010) Histone H2A C-Terminus Regulates Chromatin Dynamics, Remodeling, and Histone H1 Binding. Plos Genet, 6.

60. Arya, G. and Schlick, T. (2009) A Tale of Tails: How Histone Tails Mediate Chromatin Compaction in Different Salt and Linker Histone Environments. J Phys Chem A, 113, 4045–4059.

61. Fan, J.Y., Gordon, F., Luger, K., Hansen, J.C. and Tremethick, D.J. (2002) The essential histone variant H2A.Z regulates the equilibrium between different chromatin conformational states. Nature Structural Biology, 9, 172–176.

62. Hildreth, A.E., Ellison, M.A., Francette, A.M., Seraly, J.M., Lotka, L.M. and Arndt, K.M. (2020) The nucleosome DNA entry-exit site is important for transcription termination and prevention of pervasive transcription. Elife, 9.

63. Soboleva, T.A., Nekrasov, M., Pahwa, A., Williams, R., Huttley, G.A. and Tremethick, D.J. (2012) A unique H2A histone variant occupies the transcriptional start site of active genes. Nature Structural & Molecular Biology, 19, 25–U37.

64. Weber, C.M., Henikoff, J.G. and Henikoff, S. (2010) H2A.Z nucleosomes enriched over active genes are homotypic. Nature Structural & Molecular Biology, 17, 1500–U1136.

65. Tolstorukov, M.Y., Kharchenko, P.V., Goldman, J.A., Kingston, R.E. and Park, P.J. (2009) Comparative analysis of H2A.Z nucleosome organization in the human and yeast genomes. Genome Res, 19, 967–977.

66. Teves, S.S., Weber, C.M. and Henikoff, S. (2014) Transcribing through the nucleosome. Trends in Biochemical Sciences, 39, 577–586.

67. Rhee, Ho S., Bataille, Alain R., Zhang, L. and Pugh, B.F. (2014) Subnucleosomal Structures and Nucleosome Asymmetry across a Genome. Cell, 159, 1377–1388.

68. Chen, Y.J., Tokuda, J.M., Topping, T., Meisburger, S.P., Pabit, S.A., Gloss, L.M. and Pollack, L. (2017) Asymmetric unwrapping of nucleosomal DNA propagates asymmetric opening and dissociation of the histone core. P Natl Acad Sci USA, 114, 334–339.

69. Abraham, M.J., Murtola, T., Schulz, R., Páll, S., Smith, J.C., Hess, B. and Lindahl, E. (2015) GROMACS: High performance molecular simulations through multi-level parallelism from laptops to supercomputers. SoftwareX, 1-2, 19–25.

70. Galindo-Murillo, R., Robertson, J.C., Zgarbova, M., Sponer, J., Otyepka, M., Jurecka, P. and Cheatham, T.E. (2016) Assessing the Current State of Amber Force Field Modifications for DNA. J Chem Theory Comput, 12, 4114–4127.

71. Maier, J.A., Martinez, C., Kasavajhala, K., Wickstrom, L., Hauser, K.E. and Simmerling, C. (2015) ff14SB: Improving the Accuracy of Protein Side Chain and Backbone Parameters from ff99SB. J Chem Theory Comput, 11, 3696–3713.

72. Izadi, S., Anandakrishnan, R. and Onufriev, A.V. (2014) Building Water Models: A Different Approach. J Phys Chem Lett, 5, 3863–3871.

73. Shabane, P.S., Izadi, S. and Onufriev, A.V. (2019) General Purpose Water Model Can Improve Atomistic Simulations of Intrinsically Disordered Proteins. Journal of Chemical Theory and Computation, 15, 2620–2634.

74. Bussi, G., Donadio, D. and Parrinello, M. (2007) Canonical sampling through velocity rescaling. J Chem Phys, 126, 014101.

75. Parrinello, M. and Rahman, A. (1981) Polymorphic Transitions in Single-Crystals - a New Molecular-Dynamics Method. J Appl Phys, 52, 7182–7190.

76. Essmann, U., Perera, L., Berkowitz, M.L., Darden, T., Lee, H. and Pedersen, L.G. (1995) A Smooth Particle Mesh Ewald Method. J Chem Phys, 103, 8577–8593.

77. Hess, B., Bekker, H., Berendsen, H.J.C. and Fraaije, J.G.E.M. (1997) LINCS: A linear constraint solver for molecular simulations. J Comput Chem, 18, 1463–1472.

78. Case, D.A., Aktulga, H.M. and Belfon, K. (2020) Amber 2020, University of California, San Francisco.

